# Evidence for the cardiodepressive effects of the plasticizer di-2-ethylhexylphthalate (DEHP)

**DOI:** 10.1101/2023.05.22.541729

**Authors:** Luther M. Swift, Anysja Roberts, Jenna Pressman, Devon Guerrelli, Samuel Allen, Kazi T. Haq, Julie A Reisz, Angelo D’Alessandro, Nikki Gillum Posnack

## Abstract

Di-2-ethylhexylphthalate (DEHP) is commonly used in the manufacturing of plastic materials, including intravenous bags, blood storage bags, and medical-grade tubing. DEHP can leach from plastic medical products, which can result in inadvertent patient exposure. DEHP concentrations were measured in red blood cell (RBC) units stored between 7-42 days (23-119 μg/mL). Using these concentrations as a guide, Langendorff-perfused rat heart preparations were acutely exposed to DEHP. Sinus activity remained stable with lower doses of DEHP (25-50 μg/mL), but sinus rate declined by 43% and sinus node recovery time prolonged by 56.5% following 30-minute exposure to 100 μg/ml DEHP. DEHP exposure also exerted a negative dromotropic response, as indicated by a 69.4% longer PR interval, 108.5% longer Wenckebach cycle length, and increased incidence of atrioventricular uncoupling. Pretreatment with doxycycline partially rescued the effects of DEHP on sinus activity, but did not ameliorate the effects on atrioventricular conduction. DEHP exposure also prolonged the ventricular action potential and effective refractory period, but had no measurable effect on intracellular calcium transient duration. Follow-up studies using hiPSC-CM confirmed that DEHP slows electrical conduction in a time (15 min – 3 hours) and dose-dependent manner (10-100 μg/mL). Previous studies have suggested that phthalate toxicity is specifically attributed to metabolites of DEHP, including mono-2-ethylhexyl phthalate (MEHP). This study demonstrates that DEHP exposure also contributes to cardiac dysfunction in a dose- and time-dependent manner. Future work is warranted to investigate the impact of DEHP (and its metabolites) on human health, with special consideration for clinical procedures that employ plastic materials.

## INTRODUCTION

Phthalate plasticizer chemicals are widely used in the manufacturing of both consumer goods and medical products, including endotracheal tubes, intravenous bags, blood storage bags, and medical-grade tubing(Shelby, 2006; Ramadan *et al*., 2020). Di(2-ethylhexyl) phthalate (DEHP) is frequently used as a plasticizer additive, wherein it embeds between polyvinyl chloride polymers to soften and impart flexibility to plastic materials(DiGangi, 1999; Jaeger and Rubin, 1973). DEHP is highly susceptible to leaching because it is not covalently bound to the polyvinyl chloride matrix; phthalate leaching is increased when lipophilic solutions come into contact with plastic materials that have a large surface area, at an elevated temperature, or for a prolonged period of time(Rose *et al*., 2012; Bagel-Boithias *et al*., 2005; Karle *et al*., 1997; Loff *et al*., 2002; D’alessandro *et al*., 2016; Rock *et al*., 1986). Historically, DEHP leaching has been considered beneficial in the blood banking field, since DEHP acts as a preservative agent to reduce red blood cell (RBC) hemolysis and extend the shelf life of donated blood products(Rock *et al*., 1984; Labow *et al*., 1987; AuBuchon *et al*., 1988; Horowitz *et al*., 1985). Several studies have shown that DEHP can accumulate in stored blood products, which can result in inadvertent patient exposure(D’alessandro *et al*., 2016; Inoue *et al*., 2005; Kaestner *et al*., 2020; Huygh *et al*., 2015). Indeed, clinical exposure to phthalate chemicals has also been documented in patients undergoing multiple medical interventions that employ plastic products(Green *et al*., 2005; Weuve *et al*., 2006; Calafat *et al*., 2004; Mallow and Fox, 2014; Takahashi *et al*., 2008; Shang *et al*., 2019; Malarvannan *et al*., 2019). Accordingly, safety concerns have been raised regarding phthalate exposure and the potential negative implications on human health(Braun *et al*., 2013; Posnack, 2021; Tereshchenko and Posnack, 2019).

Epidemiological studies have reported associations between phthalate exposure and an increased risk of all-cause and cardiovascular mortality, which has prompted a renewed interest in understanding the direct effect of DEHP on the heart(Trasande *et al*., 2022; Zeng *et al*., 2022). Identifying the consequences of clinical phthalate exposure on patient outcomes is inherently difficult, given the complexity of clinical care and the heterogenous nature of patient populations. Nevertheless, experimental studies have shown that phthalate chemicals can exert cardiotoxic effects(Posnack, 2014; Gillum *et al*., 2009; Jaimes, McCullough, Siegel, Swift, McInerney, *et al*., 2019). As examples, DEHP exposure slows the spontaneous beating rate of isolated cardiomyocyte preparations(Rubin and Jaeger, 1973; Posnack *et al*., 2015), while MEHP exposure (primary metabolite of DEHP) delays atrioventricular conduction and precipitates cardiac arrest in rodents(Rock *et al*., 1987; Jaimes, McCullough, Siegel, Swift, McInerney, *et al*., 2019). Although the underlying mechanisms are incompletely understood, it has been suggested that phthalates may exert immediate effects via interaction with cholinergic receptors(Barry *et al*., 1990; Pfuderer and Francis, 1975), while longer exposure times can disrupt intercellular coupling likely via oxidative stress and/or increased matrix metalloproteinase activity(Sobarzo *et al*., 2015; Yao *et al*., 2010; Gillum *et al*., 2009). For the latter, comparative studies conducted in liver cells estimated that DEHP reduces gap junction intercellular coupling by approximately 20% within 1 hour and >80% within 16 hours of exposure(Čtveráčková *et al*., 2020).

In the current experimental study, we aimed to further understand the potential risks of phthalate exposure on cardiac physiology. We measured DEHP concentrations in stored RBC units and then used those concentrations as a guide for cardiotoxicity assessment. Specifically, we quantified the acute effects of DEHP exposure on cardiac electrophysiology using intact rodent heart preparations and human induced pluripotent stem cell-derived cardiomyocytes. Further, we investigated whether pretreatment with a cholinergic receptor antagonist (atropine) or matrix metalloproteinase inhibitor (doxycycline) could abrogate the cardiotoxic effects of DEHP.

## METHODS

### Phthalate Chemical Extraction and Quantitation

Phthalate chemical concentrations were measured in supernatant samples collected from refrigerated RBC units, stored for 7-42 days (**Figure 1**). Briefly, samples were shipped on dry ice to the University of Colorado School of Medicine Metabolomics Core. Samples were thawed on ice and phthalates were extracted from a 10 μL aliquot via vigorous vortexing for 30 min at 4°C in the presence of 5:3:2 methanol:acetonitrile:water (240 μL) containing an isotope labeled standard (1 μM DEHP ring-1,2-^13^C_2_, Cambridge Isotope Laboratories). Extraction supernatants were clarified via centrifugation at 18,000 rpm for 10 min at 4°C and analyzed immediately by ultra high-pressure liquid chromatography coupled to mass spectrometry (UHPLC-MS). Sample injection volume was 20 μL. Samples were analyzed on a Thermo Vanquish UHPLC coupled to a Thermo Q Exactive high resolution mass spectrometer. The LC was equipped with a Phenomenex Kinetex C18 column (2.1 x 150 mm, 1.7 μm) held at 45 °C (gradient information below). The mass spectrometer scanned in MS1 mode in the range of 65 to 975 m/z at a resolution of 70,000. Instrument-generated .raw files were converted to .mzXML format via RawConverter and peaks for phthalates and accompanying standards were extracted and integrated using Maven (Princeton University). Absolute concentrations were obtained using the following equation where dilution factor (DF) is 25: [Phthalate] = (Peak Area Phthalate) / (Peak Area Standard) * [Standard] * DF

**Figure 1.**
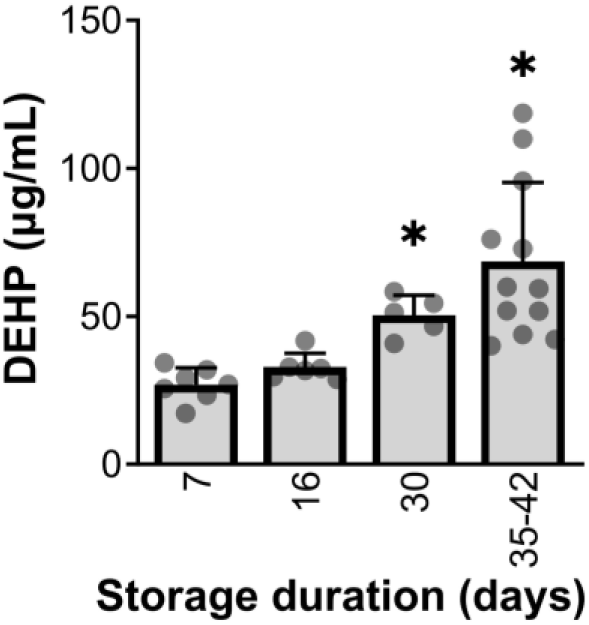
DEHP concentration increases in refrigerated RBC units with longer storage time. RBC unit supernatant samples were collected from units stored 7-42 days; DEHP levels were quantitated using ultra high-pressure liquid chromatography coupled to mass spectrometry. *One-way ANOVA with Dunnett’s multiple comparisons test. Significance compared to day 7 RBC unit, denoted by *p<0.05. n=5-12 individual units per timepoint*.

DEHP was quantified using a 5 min C18 gradient at a flow rate of 450 μL per min as previously described(Nemkov *et al*., 2019) with a single change – eluant was introduced to the MS using atmospheric pressure chemical ionization (APCI) in positive mode. APCI was performed using 10 sheath gas, 20 aux gas (both N_2_), and 350°C vaporizer temperature. The calculations of DEHP concentrations utilized the average peak area of the stable isotope labeled DEHP standard across all samples within the biological group.

### Animal model

Animal protocols were approved by the Institutional Animal Care and Use Committee within the Children’s National Research Institute and followed the Nationals Institutes of Health’s *Guide for the Care and Use of Laboratory Animals*. Experiments were performed using 2–4-month-old, mixed sex Sprague Dawley rats (strain NTac: SD, from NIH Genetic Resource stock, Taconic Biosciences: Germantown New York). Animals were housed in conventional acrylic cages in the Research Animal Facility, under standard environmental conditions (12:12 hours light:dark cycle, 18-25° C, 30-70% humidity).

### Isolated heart preparation and Langendorff perfusion

Animals were anesthetized with isoflurane (2-4%) for approximately 10 minutes until a surgical depth of anesthesia was reached, as determined by a tail pinch test. Animals were then euthanized by exsanguination following heart excision. Briefly, the heart was excised and placed in ice-cold cardioplegia; thereafter, the aorta was cannulated and the intact heart was transferred to a temperature controlled (37°C), constant pressure (70 mm/Hg) Langendorff-perfusion system. The heart was retrograde perfused with a modified Krebs-Henseleit buffer (mM: 118.0 NaCl, 3.3 KCl, 1.2 MgSO_4_, 1.2 KH_2_PO_4_, 24.0 NaHCO_3_, 10.0 glucose, 2.0 sodium pyruvate, 10.0 HEPES buffer, 2.0 CaCl_2_) bubbled with carbogen (95% O_2_, 5% CO_2_). Intact heart preparations were allowed to equilibrate for 10-15 minutes on the perfusion system; once the myocardium was stabilized, blebbistatin was added to the perfusate to reduce metabolic demand and suppress contractile motion(Fedorov *et al*., 2007; Swift *et al*., 2021). A low dose of blebbistatin (1.25 μg/mL) was used for electrophysiology studies, while a slightly higher dose of blebbistatin (3.75 μg/ml) was used for imaging studies(Swift *et al*., 2012). To take into account inter-animal variability, each animal served as its own control with baseline electrophysiology recordings collected before treatment. Time control experiments were also performed to account for any slight deviation in myocardial physiology that might occur during prolonged Langendorff-perfusion (150 minutes). DEHP experiments were conducted by diluting the neat stock liquid into 0.1% DMSO (to aid in solubility), which was diluted in Krebs-Henseleit buffer to reach a final circulating concentration of 25-100 μg/mL. This concentration was selected to mimic a clinical exposure to DEHP following contact with plastic medical products, similar to exposure via blood transfusion(Rael *et al*., 2009; Sjoberg *et al*., 1985). In a subset of studies, intact heart preparations were pretreated with either 1 μM atropine (muscarinic antagonist) or 20 μM doxycycline (matrix metalloproteinase inhibitor) – concentrations that have previously been shown to augment the effects of acetylcholine and matrix metalloproteinases(Hoover and Neely, 1997; Scannevin *et al*., 2017).

### Electrophysiology measurements

*Ex vivo* electrocardiograms (ECG) were recorded continuously from excised, intact heart preparations by placing needle electrodes within the perfusate bath. ECGs were recorded using a PowerLab acquisition system, in conjunction with a differential amplifier (Warner Instruments, Holliston MA) and a 60 Hz noise eliminator (Hum Bug: Digitimer, Fort Lauderdale FL). From each ECG recording, the heart rate, P-wave duration, and PR interval were measured during sinus rhythm. Coronary flow rate was monitored continuously using an inline flow sensor (Transonic, Ithaca NY) positioned above the aortic cannula. In a subset of studies, microelectrode arrays (MEA, MappingLabs Ltd., United Kingdom) were positioned in two configurations to measure activation time: one MEA on the right atrium and the second MEA positioned on either the left atrium (to measure atrial conduction) or the left ventricle (to measure atrioventricular conduction). Electrical signals were acquired with a multi-channel amplifier using eMapRecord software and analyzed using EMapScope (MappingLabs Ltd.).

Programmed electrical stimulation was implemented by placing two small stimulation electrodes (Harvard Biosciences, Holliston MA) on the right atrium and left ventricle. An electrophysiology stimulator (Bloom: Fisher Medical, Wheat Ridge CO) was set to 1 ms pulse width at 1.5x the minimum threshold current (∼0.7-1.2 mA)(Swift *et al*., 2019). To assess sinus node recovery time (SNRT), spontaneous sinus activity was suppressed via fast atrial pacing (20 beats, S1-S1) and SNRT was measured as the time delay between the last paced beat and the first spontaneous P-wave. Corrected SNRT (cSNRT) was measured as the difference between SNRT and the RR-interval during sinus rhythm. To identify the Wenckebach cycle length (WBCL), atrial pacing was initiated (S1-S1) and the pacing cycle length (PCL) was decremented until atrioventricular (AV) conduction failed. WBCL was defined as the shortest S1-S1 interval that resulted in 1:1 AV conduction, and 2:1 capture was defined as the longest S1-S1 interval that resulted in 2:1 capture of AV conduction. The AV node effective refractory period (AVNERP) was determined by introducing an extrastimulus (atria pacing) and pinpointing the shortest S1-S2 interval that resulted in 1:1 AV conduction. The ventricular effective refractory period (VERP) was determined by decrementing the S1-S2 interval (ventricular pacing) to identify the shortest S1-S2 interval that resulted in ventricular depolarization.

### Fluorescence imaging and analysis

Isolated hearts were loaded with a calcium indicator dye (100 μg Rhod-2, AM; AAT Bioquest, Sunnyvale CA) and/or a potentiometric dye (62 μg RH237; AAT Bioquest). Fluorescence signals were acquired by illuminating the epicardial surface with an LED light with excitation filter (535 ± 25 nm; ThorLabs, Sterling VA); emission filters were used to collect the calcium signal (585 ± 20 nm) or voltage signal (>710 nm)(Jaimes, McCullough, Siegel, Swift, Hiebert, *et al*., 2019). Images were collected using high-speed cameras at 400-600 fps (Dimax CS4: PCO-Tech, Kelheim Germany; Optical Mapping System from MappingLabs Ltd equipped with Prime BSI cameras, Teledyne Photometrix). Optical action potentials and calcium transients were measured at the apex of the left ventricle; fluorescence signals were analyzed using custom software (*CaMat*(Jaimes *et al*., 2016), *Kairosight*(Cooper *et al*., 2021; Haq *et al*., 2023)).

### Human induced pluripotent stem cell-derived cardiomyocyte (hiPSC-CM) recordings

Cryopreserved hiPSC-CMs (iCell cardiomyocytes^2^, female donor #01434, Fujifilm) were thawed and plated onto fibronectin-coated microelectrode arrays (MEA) at a density of 40-65,000 cells/well (Axion BioSystems, Atlanta GA). Cells were defrosted in iCell cardiomyocyte plating media in a cell culture incubator (37°C, 5% CO_2_) for 2 hours, thereafter cells were cultured in iCell maintenance media (Fujifilm). hiPSC-CM formed a confluent monolayer within three days after plating and were used thereafter for conduction velocity measurements. Briefly, hiPSC-CM were equilibrated in a temperature (37°C) and gas controlled (5% CO_2_) live cell monitoring system (Maestro Edge, Axion) for 20-minutes. Electrical stimulation was applied (1.5 Hz frequency, 800 mV, 20 μA, 400 μs stimulus duration) and conduction velocity was measured across a 16-electrode array under baseline conditions and again after chemical exposure (0.1% DMSO, 10-100 μg/mL DEHP). In a subset of studies, hiPSC-CM were pretreated with either 1 μM atropine (antimuscarinic) or 20 μM doxycycline (matrix metalloproteinase inhibitor) before DEHP exposure.

### Statistical analysis

Results are reported as the mean ± standard deviation. Normality was assessed for each data set (Shapiro-Wilk test). Two group comparisons (e.g., baseline vs 60-minute control) were analyzed using a two-tailed T-test. Three or more group comparisons (e.g., dose and time-course studies) were analyzed using a one-way ANOVA test with Dunnett’s (parametric data sets) or Dunn’s (nonparametric) multiple comparisons test. Comparisons with two independent factors (e.g., APD measurements across multiple pacing frequencies) were analyzed using two-way ANOVA with Holm-Sidak test for multiple comparisons. Significance is denoted in each figure with an asterisk (p-value <0.05).

## RESULTS

### DEHP exerts a negative chronotropic effect in a dose-dependent manner

Phthalate plasticizers have been shown to leach from medical plastic devices, resulting in unintentional patient exposure. Prior studies demonstrated that DEHP can accumulate in stored blood or migrate from polyvinyl chloride tubing, reaching concentrations as high as 400 μg/mL(Rael *et al*., 2009; Loff *et al*., 2002; Karle *et al*., 1997; Mallow and Fox, 2014). In agreement, we found that DEHP concentrations increased in RBC units during refrigerated storage, beginning at 17-34 μg/mL in fresh units (7 days storage) and increasing to 40-119 μg/mL in units near expiry (35-42 days; p<0.05; **Figure 1**). Notably, the Recipient Epidemiology and Donor Evaluation Strategy (REDS)-III found that 9-21% of transfused RBC units are stored for at least 35 days(Glynn *et al*., 2016). Using these concentrations as a guide, we next performed dose-response studies to measure the direct effect of DEHP exposure on cardiac function using Langendorff-perfused heart preparations (**Figure 2**). For these initial studies, we focused on cardiac automaticity as a primary endpoint, since prior work has shown that DEHP slows the beating rate of isolated cardiomyocytes(Rubin and Jaeger, 1973; Posnack *et al*., 2015). Sinus rate remained stable under control conditions, declining by only 3.5% during prolonged time course studies (baseline: 322±43, time control (150 min): 311±38 BPM, p=0.6, **Figure 2A**). Sinus rate also remained stable following an acute 30-minute exposure to lower doses of DEHP (baseline: 292±26; 25 μg/mL DEHP: 299±41; 50 mg/mL DEHP: 272±52 BPM; **Figure 2A**). However, a negative chronotropic effect was observed soon after increasing the circulating concentration to 100 μg/mL DEHP, which decreased the sinus rate by 43% within 30-minutes (baseline: 292±26, 100 μg/mL DEHP: 167±66 BPM, p<0.001). Only minor variations in sinus rhythm were observed under control conditions (baseline RMSSD: 1.13±0.75, time control (150 min): 1.86±1.31 ms, p=0.1) and following acute exposure to a low concentration of DEHP (baseline RMSSD: 1.12±0.64; 25 μg/mL DEHP: 2.91±3.0 ms; **Figure 2A**). More extreme fluctuations and abnormal sinus rhythms were observed after increasing the circulating concentration to 50 μg/mL (RMSSD: 8.3±9.7 ms, p<0.01) or 100 μg/mL DEHP (RMSSD: 22.6±22.5 ms, p<0.001).

**Figure 2.**
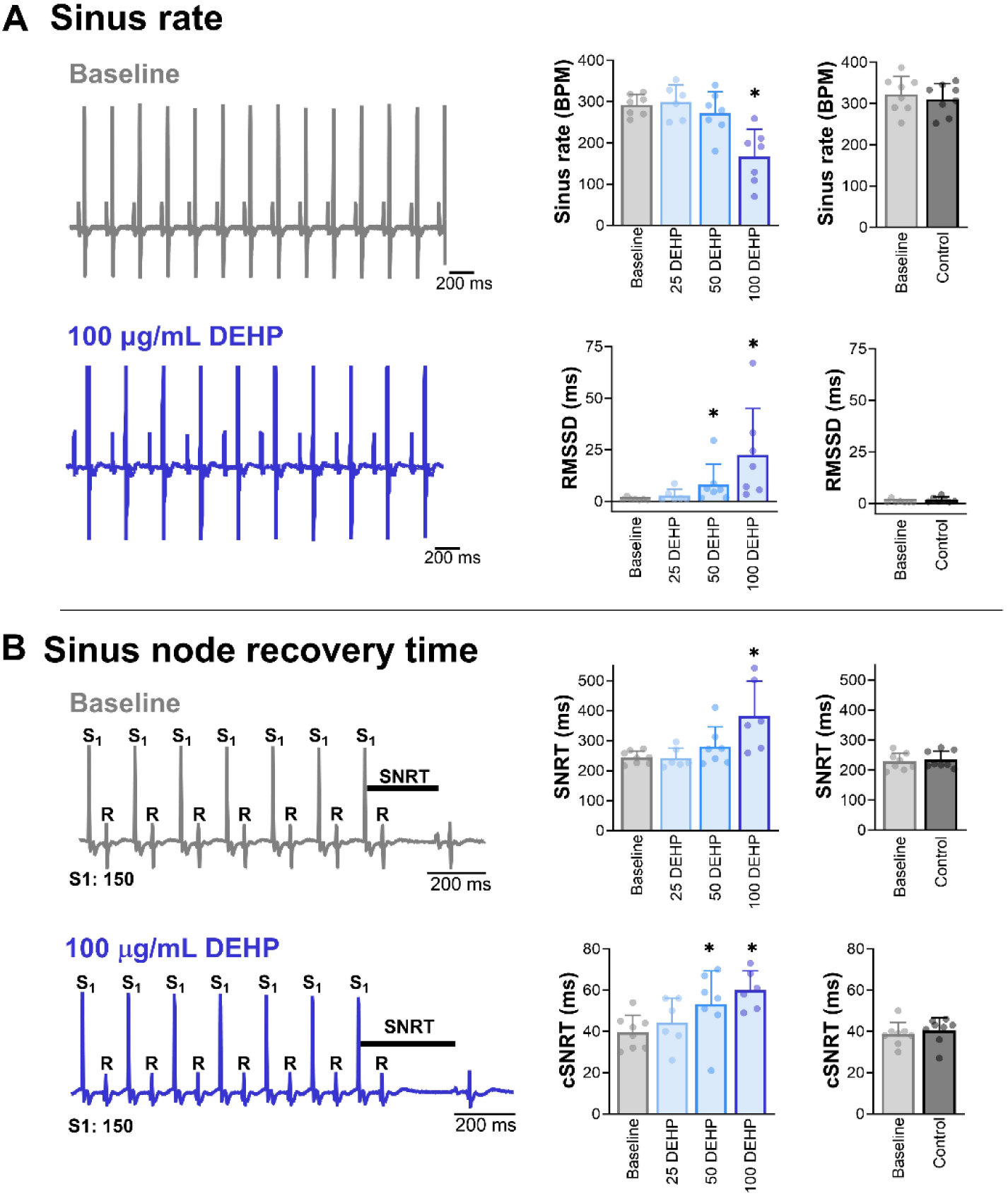
DEHP slows sinus rate and recovery time in a dose-dependent manner. **A)** <u>Left:</u> Pseudo-ECG recordings from Langendorff-perfused hearts at baseline and after exposure to 100 μg/mL DEHP (30 min). <u>Middle</u>: Sinus rate slows and sinus rhythm becomes more erratic following acute exposure to higher concentrations of DEHP (30 min exposure each dose, 25-100 μg/mL DEHP). <u>Right</u>: Time-control experiments show that sinus rate and rhythm were comparable at baseline and after prolonged perfusion (150 min) with control media. **B)** <u>Left</u>: Measurements of sinus node recovery time (SNRT) at baseline and after exposure to 100 μg/mL DEHP (30 min). <u>Middle</u>: DEHP exposure slows sinus node recovery. <u>Right</u>: Sinus node recovery remains unchanged in time-controlled experiments with control media perfusion (150 min). *Cardiac preparations were omitted from analysis if external pacing with 1:1 capture could not be achieved. Comparisons were analyzed by two-tailed T-test (baseline vs control) or one-way ANOVA (dose response) with multiple comparisons test. Significance compared to baseline denoted by *p<0.05. n≥6 individual heart preparations per measurement.* RMSSD = root mean square of successive differences between beats, SNRT: sinus node recovery time, cSNRT = heart rate corrected SNRT

We also quantified sinus node recovery time (SNRT), a measurement of automaticity that is evaluated using overdrive pacing. SNRT remained stable under control conditions (baseline: 229±28; time control (150 min): 235±28 ms) and following an acute 30-minute exposure to lower doses of DEHP (baseline: 244±21; 25 μg/mL DEHP: 242±33; 50 μg/mL DEHP: 280±66 ms; **Figure 2B**). While acute exposure to 100 μg/mL DEHP significantly impaired sinus function, as demonstrated by a longer SNRT (baseline: 244±21; 100 μg/mL DEHP: 382±117 ms, p<0.05). Similar results were obtained when SNRT measurements were corrected for heart rate, wherein the baseline measurement (cSNRT: 39.6±8.2) was significantly shorter than DEHP-exposed hearts (50 μg/mL DEHP: 53.1±16.2; 100 μg/mL DEHP: 60.0±9.4 ms, p<0.05; **Figure 2B**). Further, in one-quarter of the high dose studies (2 of 8 experiments), SNRT could not be determined as atrial pacing either failed to elicit a ventricular response or sinus activity was completely absent as discerned by the lack of a p-wave.

### DEHP exerts a negative chronotropic effect in a time-dependent manner

Prolonged exposure to phthalates can occur in the clinical setting over multiple hours, as is the case in patients receiving a blood transfusion, exchange transfusion, circulatory support (cardiopulmonary bypass, extracorporeal membrane oxygenation), or those receiving intensive care(Ramadan *et al*., 2020). Accordingly, we investigated the time-dependent effects of low (25 μg/mL) and high (100 μg/mL) DEHP exposure on cardiac electrophysiology. In this subset of studies, 25 μg/mL DEHP had no effect on the spontaneous beating rate, rhythm, sinus node function, or flow rate as measured in Langendorff-perfused hearts (**Figure 3B-E**). However, exposure to 100 μg/mL DEHP negatively impact cardiac electrophysiology in a time-dependent manner. We observed a progressive decline in the beating rate by 27.3% at 60 minutes (baseline: 326.3±40.8, DEHP 60 min: 237.1±47.7 BPM, p<0.0001) and 46.7% at 90 minutes (DEHP 90 min: 173.6±99.3 BPM, p<0.0001; **Figure 3A,B**). Atrioventricular uncoupling was more commonly observed with a longer exposure time, which resulted in divergent beating rates between the atria and ventricles. As such, the ventricular rate declined, on average, by 55.6% at 90 minutes exposure (baseline: 326.3±40.8, DEHP 90 min: 144.8±89.5 BPM, p<0.0001; **Figure 3B**). Prolonged exposure time also impacted the ventricular rhythm, as denoted by a marked increase in beat rate variability (baseline RMSSD: 0.86±0.2, DEHP 60 min: 17.7±14.8, DEHP 90 min: 83.6±71.3 ms, p<0.0001; **Figure 3C**). SNRT was also delayed with a longer exposure time, increasing by 52% at 60 minutes and 111% at 90 minutes (baseline: 233.1±28.9, DEHP 60 min: 353.6±147.1, DEHP 90 min: 492.6±329.6 ms, p<0.001; **Figure 3D**). A similar delay was observed when SNRT was corrected for sinus rate (cSNRT), with recovery time prolonged by 103% at 60 minutes and 387% at 90 minutes (baseline: 36.6±9.6, DEHP 60 min: 75.8±59.6, DEHP 90 min: 178.2±228.4 ms, p<0.01). SNRT and cSNRT became more difficult to measure with prolonged exposure, as atrial pacing either failed to elicit a ventricular response or sinus activity did not recover after pacing. Finally, it was observed that coronary flow dropped precipitously during the course of each DEHP exposure study, decreasing by 56% at 60 minutes and 66% at 90 minutes (baseline: 5.7±1.3, DEHP 60 min: 2.5±1.7, DEHP 90 min: 1.9±1.7 ml/min, p<0.0001; **Figure 3E**).

**Figure 3.**
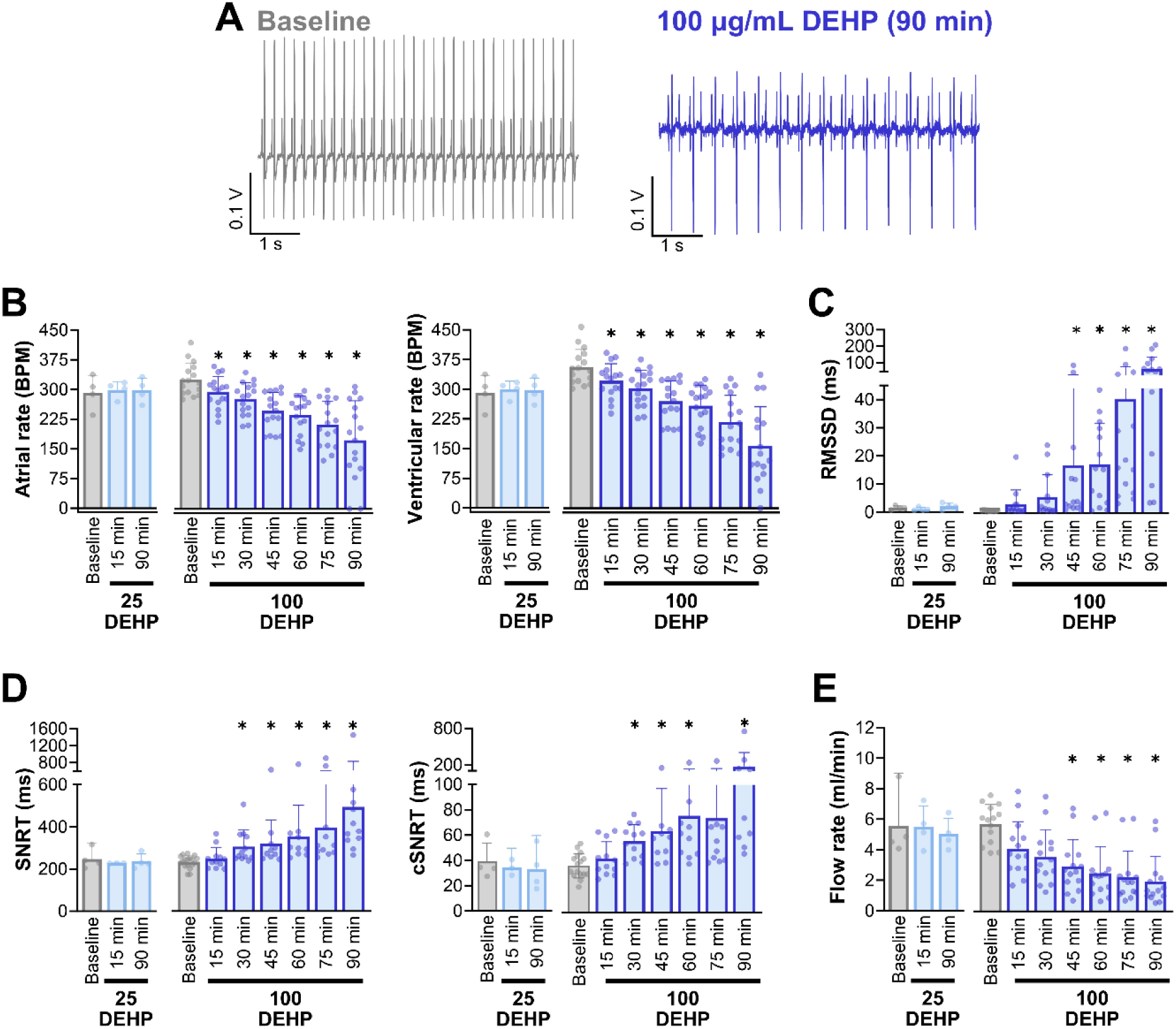
DEHP slows sinus rate and recovery time in a time-dependent manner. **A)** Pseudo-ECG recordings from Langendorff-perfused hearts at baseline and after 90-minute exposure to 100 μg/mL DEHP (note atrioventricular uncoupling). Low dose DEHP (25 μg/mL) had no measurable effect on cardiac electrophysiology; prolonged exposure to high dose DEHP (100 μg/mL) resulted in **B)** slower atrial and ventricular beat rate, **C)** increased beat rate variability, **D,E)** longer sinus node recovery time, and **F)** reduced coronary flow. *Cardiac preparations were omitted from analysis if external pacing with 1:1 capture could not be achieved. Comparisons were analyzed by one-way ANOVA with multiple comparisons test. Significance compared to baseline denoted by *p<0.05. n≥4 individual heart preparations per measurement.* RMSSD = root mean square of successive differences between beats, SNRT: sinus node recovery time, cSNRT = heart rate corrected SNRT

### DEHP exposure delays inter-atrial and atrioventricular (AV) conduction

We previously reported that DEHP exposure slows conduction velocity across cardiomyocyte monolayers(Gillum *et al*., 2009). Herein, we investigated the impact of DEHP on electrical conduction in intact heart preparations, a more complex model that preserves the cardiac conduction system and three-dimensional cell-cell interactions. Even with prolonged exposure (90 minutes), 25 μg/mL DEHP exposure had no measurable effect on atrioventricular (AV) conduction (**Figure 4**). Yet, AV conduction slowing was prominent in the presence of 100 μg/mL DEHP. During sinus rhythm, the PR interval lengthened by 23.7% after 15-minutes exposure to DEHP (baseline: 39.3±7.5, 100 μg/mL DEHP: 48.6±15.9 ms, p<0.05) and 69.4% after 60-minutes DEHP exposure (100 μg/mL DEHP: 66.6±19.3 ms, p<0.0001). With prolonged DEHP exposure, PR interval measurements became inconsistent, as AV conduction delay (1^st^ degree heart block) progressed to 2^nd^ and 3^rd^ degree heart block in 64% of heart preparations at 60-minutes (**Figure 4A**). Accordingly, PR interval measurements were only reported for coupled beats.

**Figure 4.**
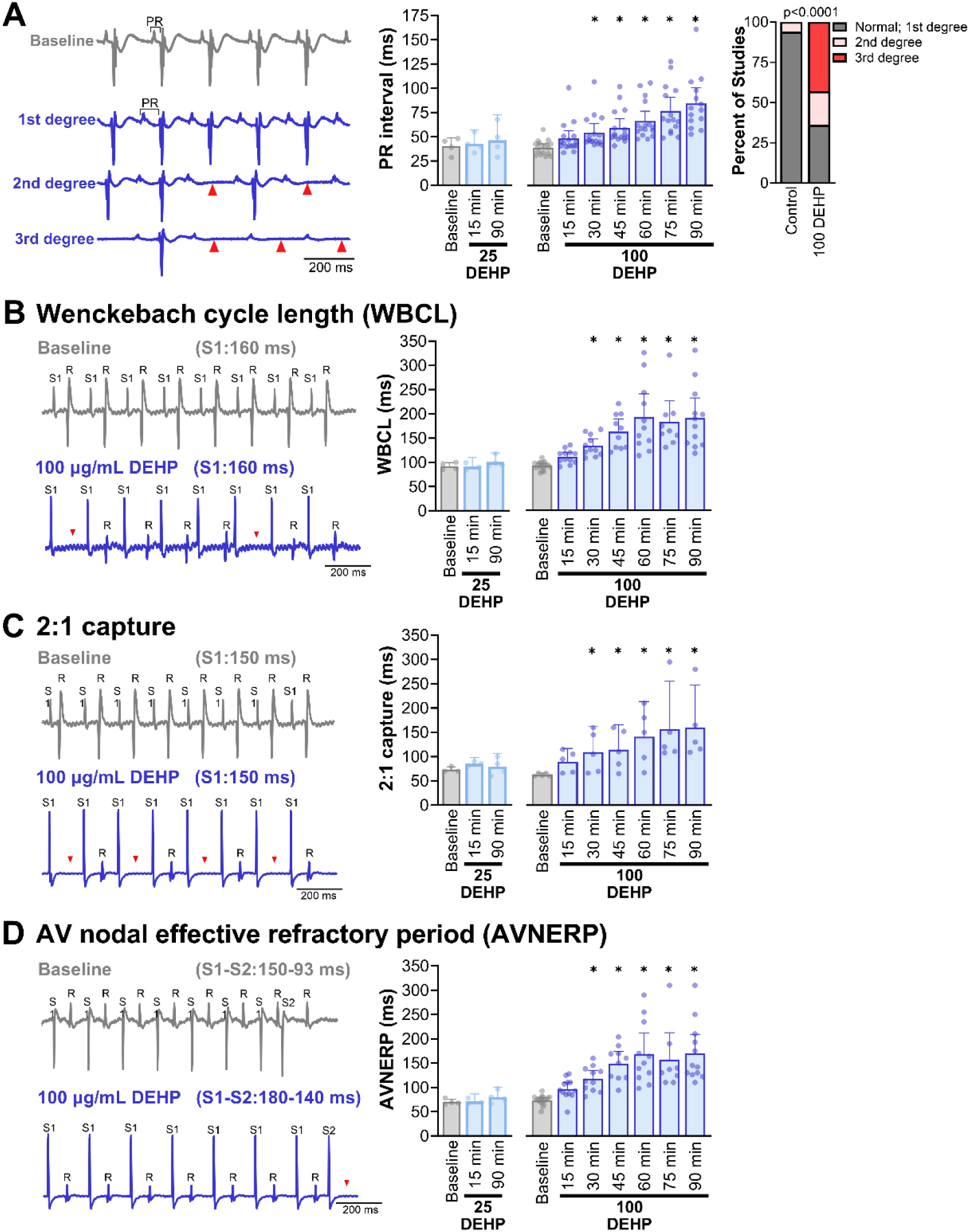
DEHP exposure increases atrioventricular conduction time and refractory period. **A)** High dose DEHP (100 μg/mL) exposure slows atrioventricular conduction in a time-dependent manner, first observed as lengthening of the PR interval (1^st^ degree heart block), then occasional dropped beats that fail to propagate to the ventricles (2^nd^ degree), and then complete 3^rd^ degree atrioventricular block with dissociation between atrial and ventricular signals. **B)** Programmed electrical stimulation at the atria shows a progressive increase in WBCL, **C)** 2:1 (atria:ventricular) capture at slower pacing frequencies, and **D)** increased AV node refractoriness at slower pacing frequencies at high dose DEHP (100 μg/mL). *Cardiac preparations were omitted from analysis if external pacing could not be achieved. Comparisons were analyzed by one-way ANOVA with multiple comparisons test. Significance compared to baseline denoted by *p<0.05. n≥4 individual heart preparations per measurement*.

Programmed electrical stimulation was used to further interrogate the effects of DEHP on AV nodal function (**Figure 4B-D**). Wenckebach cycle length (WBCL) was identified by dynamically pacing the right atrium at an increasing rate, until ventricular conduction failed. Following high dose DEHP (100 μg/mL) exposure, AV conduction failure occurred more readily at slower pacing frequencies (longer pacing cycle lengths), as indicated by a longer WBCL. For example, WBCL increased by 44.8% after 30-minutes and 108.5% after 60-minutes exposure to 100 μg/mL DEHP (baseline: 92.2±8.2, 30-min: 133.5±21.2, 60-min: 192.2±71.5 ms cycle length, p<0.001; **Figure 4B**). In a similar manner, dynamic pacing was applied to the right atrium at an increasing rate until 2:1 AV conduction block occurred. Following high dose DEHP (100 μg/mL) exposure, 2:1 AV block occurred at slower pacing frequencies within ∼45 minutes (baseline: 62.6±3.1, 45-min: 114.4±41.3 ms cycle length, p<0.05; **Figure 4C**). To determine the AV node effective refractory period (AVNERP), an extrastimulus pacing protocol was applied to the right atrium, wherein the cycle length of the last stimulus (S2) was progressively shortened until ventricular conduction failed. At baseline, AVNERP was reached at an S1-S2 interval of 150 ms (S1) and 73.8±9.5 ms cycle length (S2), which increased to 117.5±25.4 ms cycle length (S2) after 30-minutes exposure to 100 μg/mL DEHP. With longer exposure times, external pacing failed to stimulate the ventricles until exceedingly slow intervals were used (90-min: 170.3±60.8 ms cycle length (S2), p<0.0001; **Figure 4D**).

Using MEAs positioned on the surface of the heart, we also measured a three-fold increase in inter-atrial activation time (baseline: 7.2±3.7, 100 μg/mL DEHP 60 min: 21.3±4.6 ms, p<0.05), and a two-fold increase in AV activation time (baseline: 37.6±9.6, 100 μg/mL DEHP 60 min: 85.2±35.7 ms; p<0.005; **Figure 5B-C**). Delayed AV conduction was readily observed using optical mapping techniques (**Figure 5A**); in the presented example, atrial activation precedes ventricular activation by ∼40 ms at baseline and ∼80 ms after DEHP exposure (100 μg/mL, 60 minutes).

**Figure 5.**
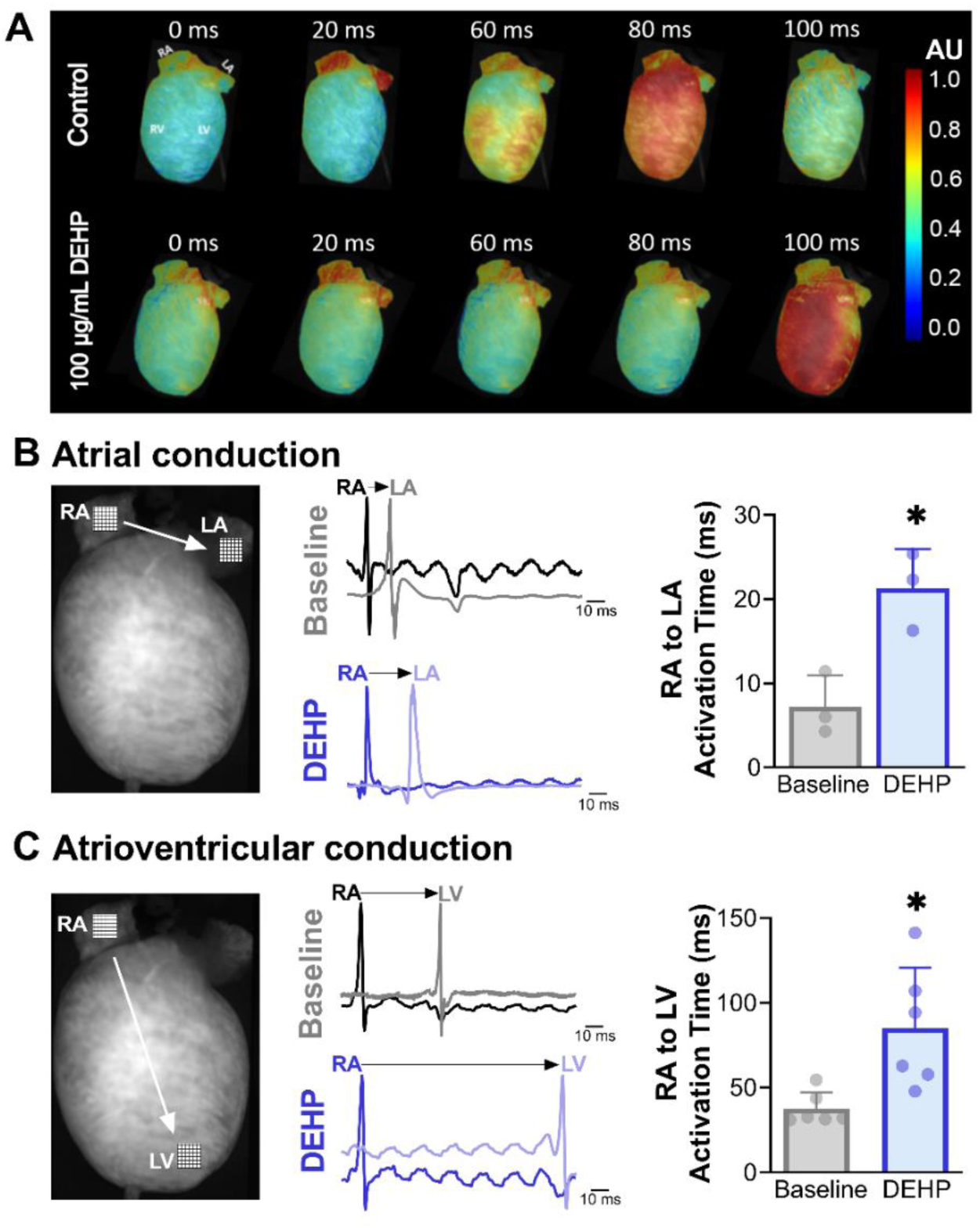
DEHP exposure delays inter-atrial and atrioventricular conduction. **A)** Time series images show the activation time (red) progressing from the atria to the ventricles, during sinus rhythm, in control and 100 μg/mL DEHP-treated heart preparations (60 minutes). **B)** Inter-atrial activation time measured using two microelectrode arrays positioned on the right atria (RA) and left atria (LA). **C)** Atrioventricular activation time measured using two microelectrode arrays positioned on the right atria (RA) and left ventricle (LV). Note the delay in activation time after 100 μg/mL DEHP exposure. AU: arbitrary units. *Comparisons were analyzed by two-tailed T-test (baseline vs DEHP); significance denoted by *p<0.05. n≥3 individual heart preparations*.

### DEHP exposure slows epicardial conduction velocity and prolongs action potential duration

Our lab, and others, have previously reported that DEHP disrupts gap junction intercellular communication in cell preparations(Gillum *et al*., 2009; Di Lorenzo *et al*., 2020; Čtveráčková *et al*., 2020). Reduced gap junction intercellular communication can slow both atrioventricular conduction velocity (described earlier) and epicardial conduction velocity. Using a voltage-sensitive dye and high-speed optical mapping, we observed that apicobasal conduction velocity remained stable in heart preparations perfused with control media, between baseline (64.8±10.3 cm/sec) and 60-minutes perfusion (64.1±14.9 cm/sec; **Figure 6A**). While apicobasal conduction velocity slowed by 15.6% between baseline (58.9±5.8 cm/sec) and 60-minutes perfusion with 100 μg/ml DEHP (49.7±5.5 cm/sec). Optical mapping of transmembrane voltage signals was also performed to evaluate ventricular action potential duration (APD), using a dynamic pacing protocol (S1-S1). Relative to time-matched controls (60-min perfusion), 100 μg/mL DEHP exposure prolonged the ventricular APD at 30% and 80% repolarization at both slower and faster pacing frequencies (**Figure 6B, D, E**). As an example, at 200 PCL DEHP exposure increased APD_30_ by 29.9% (control: 18.4±2.3, 100 μg/mL DEHP: 23.9±3.5 ms) and APD_80_ by 29.4% (control: 54.1±5.8, 100 μg/mL DEHP: 70.0±5.6 ms). Using an extrastimulus pacing protocol (S1-S2), we also observed that 60-minutes perfusion with 100 μg/mL DEHP lengthened the ventricular effective refractory period (55.9±11.1 ms) as compared to time-matched controls (40.5±7 ms; **Figure 6C**). Finally, we monitored the effects of DEHP exposure on intracellular calcium handling, but did not detect any measurable difference in the ventricular calcium transient duration time at 30% or 80%. At 200 PCL, CaD_30_ (control: 51.4±7 ms, 100 μg/mL DEHP: 53.1±5.5 ms) and CaD_80_ (control: 92.5±7.1 ms, 100 μg/mL DEHP: 100.0±8.9 ms) were comparable after 60-minutes perfusion (**Figure 6F, G**).

**Figure 6.**
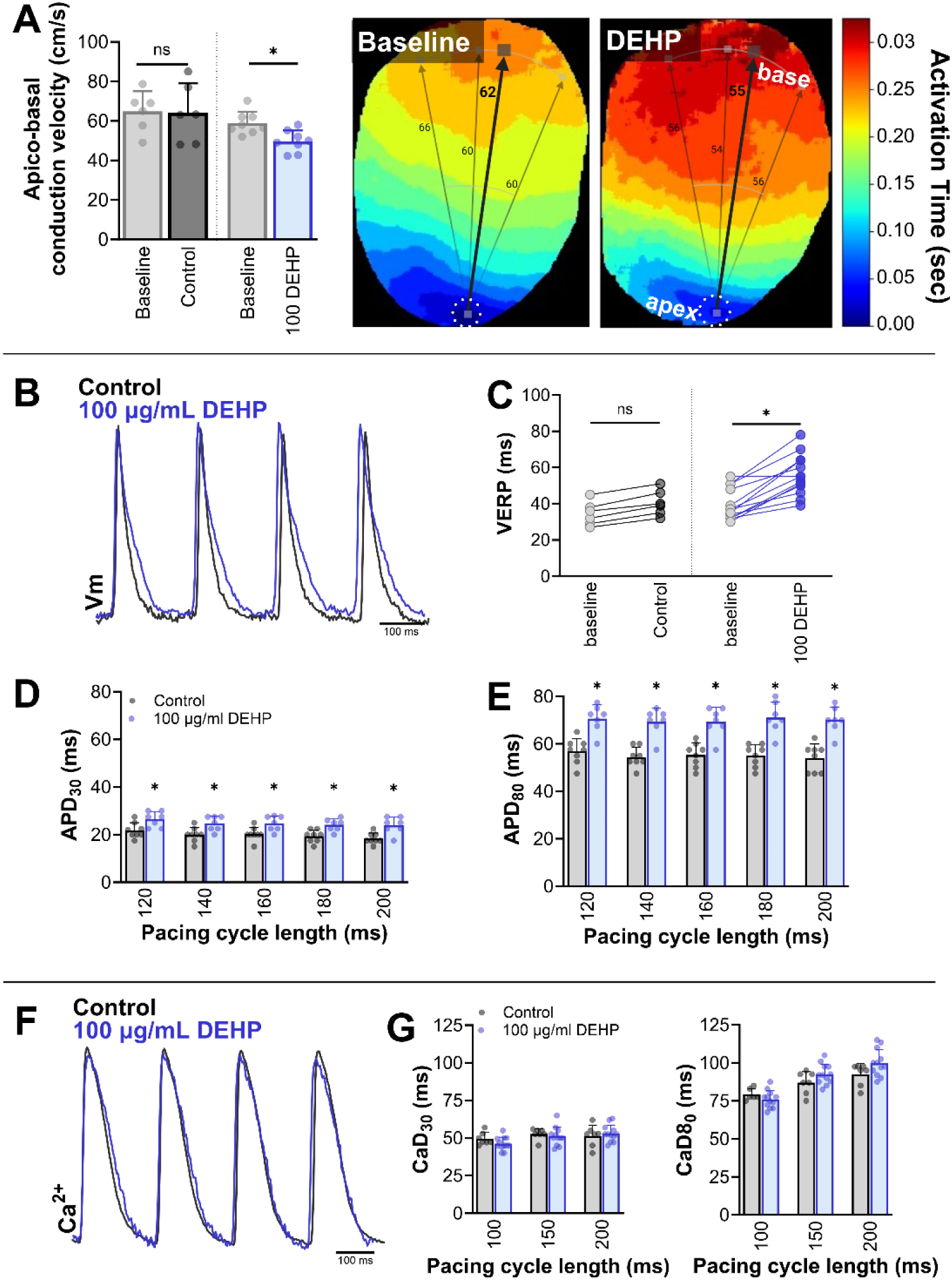
DEHP exposure slows epicardial conduction velocity and prolongs action potential duration. **A)** 100 μg/mL DEHP exposure slows apico-basal conduction velocity, compared to time-matched controls. **B-E)** 60-minutes DEHP exposure lengthens the ventricular action potential duration at 30% (APD_30_) and 80% repolarization (APD_80_) and ventricular effective refractory period (VERP). **F)** DEHP exposure had no measurable effect on the calcium transient duration time at 30% (CaD_30_) or 80% (CaD_80_). *Cardiac preparations were omitted from analysis if external pacing could not be achieved, or poor signal:noise prevented accurate measurements. VERP comparisons were analyzed by two-tailed T-test. Action potential and calcium transient comparisons were analyzed by two-way ANOVA with Holm-Sidak test for multiple comparisons. Significance between time-matched controls and DEHP (60-minutes) denoted by *p<0.05. n≥6 individual heart preparations per measurement*.

### Effects of DEHP on cardiac electrophysiology are not abolished by muscarinic antagonism

Previous studies implicated cardiac muscarinic receptors as a potential target for phthalate plasticizers, since atropine-treatment blunted the cardiodepressive effects of phthalates in both goldfish and human atrial trabecular preparations(Barry *et al*., 1990; Pfuderer and Francis, 1975). We tested this possible mechanism by pretreating Langendorff-perfused heart preparations for 10-15 minutes with 1 μM atropine, a concentration that has been shown to augment the effects of acetylcholine(Hoover and Neely, 1997), followed by exposure to 100 μg/mL DEHP. Across all the tested variables, the effects of DEHP on sinus activity and atrioventricular conduction persisted in the presence of atropine (**Figure 7A-F**). In this set of studies, ventricular rate slowed by 51.5% and SNRT increased by 123.5% in hearts with 100 μg/mL DEHP plus atropine, versus atropine alone.

**Figure 7.**
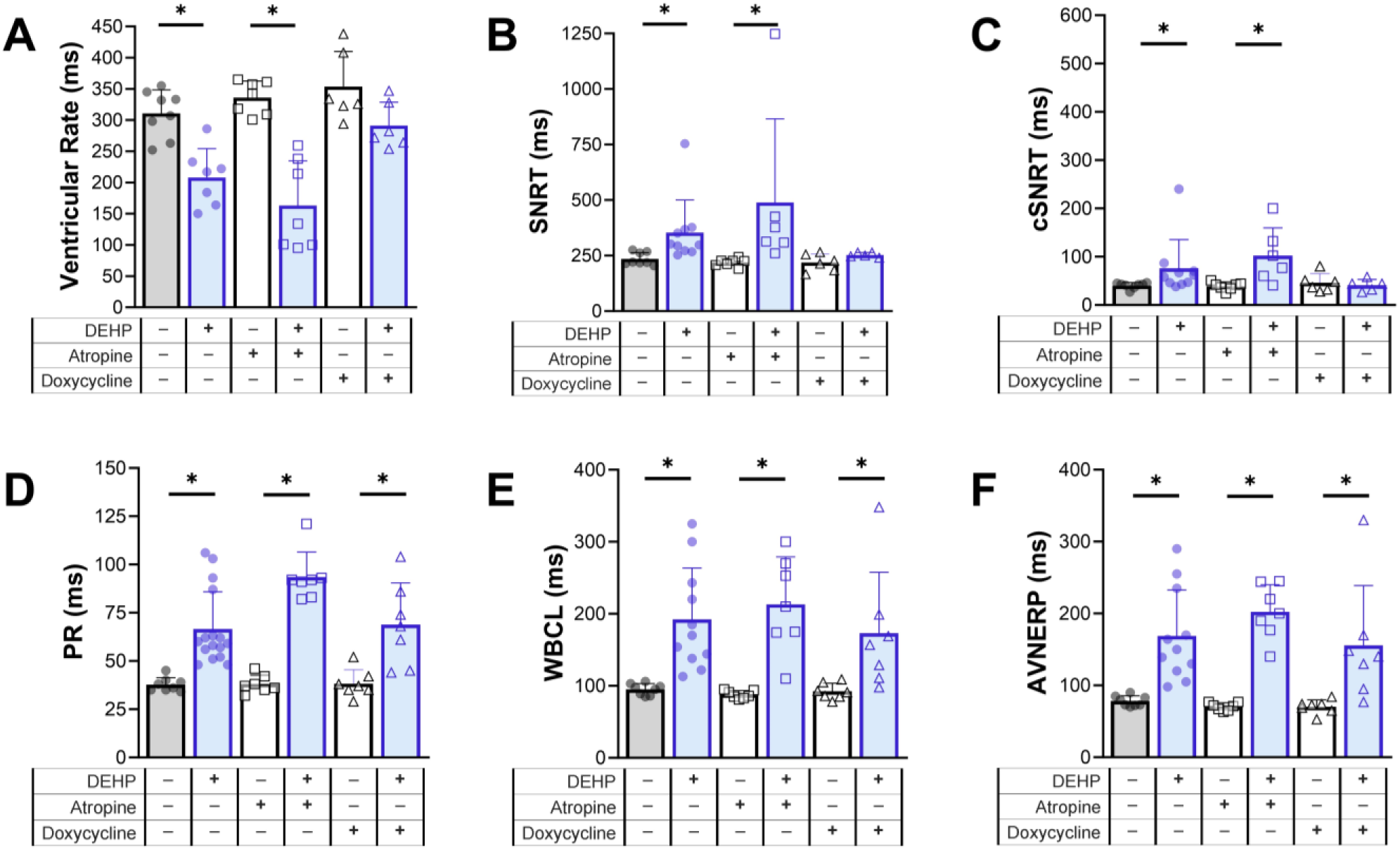
Effects of DEHP are partially relieved by inhibition of matrix metalloproteinases but not muscarinic antagonism. Langendorff-perfused hearts were exposed to 100 μg/mL DEHP for 60-minutes, without or with pretreatment using atropine (muscarinic antagonist) or doxycycline (matrix metalloproteinase inhibitor). Cardiac parameters were assessed relative to the vehicle (control, atropine, or doxycycline alone), including: **A)** ventricular beating rate, **B)** sinus node recovery time (SNRT), **C)** heart rate corrected SNRT, **D)** PR interval, **E)** Wenckebach cycle length (WBCL), **F)** atrioventricular nodal effective refractory period (AVNERP). *Cardiac preparations were omitted from analysis if external pacing could not be achieved. Comparisons to respective vehicle were analyzed by two-tailed T-test. Significance denoted by *p<0.05. n≥5 individual heart preparations per measurement*.

### Effects of DEHP are partially relieved by inhibition of matrix metalloproteinases

Previous work indicates that DEHP can disrupt intercellular coupling(Gillum *et al*., 2009; Zhao *et al*., 2022), and this effect may be mechanistically linked to an upregulation of matrix metalloproteinase activity(Yao *et al*., 2010; Posnack *et al*., 2011) that modulates gap junction proteins and influences the electrical properties of the heart(Lindsey *et al*., 2006; Wang *et al*., 2014). We tested this possible mechanism by pretreating heart preparations for 10-15 minutes with 20 μM doxycycline, a synthetic matrix metalloproteinase inhibitor(Scannevin *et al*., 2017). Doxycycline (doxy) reduced the depressive effects of DEHP on sinus activity, as demonstrated by SNRT (Doxy alone: 218.5±43.3, DEHP+doxy: 253.3±10.1 ms, p=0.4), cSNRT (doxy alone: 41±12.1 ms, DEHP+doxy: 46±18.9 ms, p=0.7), and beating rate measurements (Doxy alone: 353.8±56.1, DEHP+doxy: 291.2±37.5 BPM, p=0.051; **Figure 7A-F**). However, doxycycline pretreatment did not ameliorate the effects of DEHP on atrioventricular conduction, as the PR interval lengthened by 79.8%, WBCL increased by 87.2%, and AVNERP increased by 121.5% in hearts with 100 μg/mL DEHP plus doxycycline, versus doxycycline alone.

### DEHP slows cardiac conduction velocity in hiPSC-CM monolayers

DEHP has been shown to reduce gap junction intercellular communication, but this effect may be species-specific in select tissues (e.g., liver)(Isenberg *et al*., 2000; Kamendulis *et al*., 2002). To test the applicability of our findings on human cardiomyocytes, we measured conduction velocity using hiPSC-CM monolayers plated atop a microelectrode array (paced at 1.5 Hz). DEHP-treatment slowed conduction velocity in a dose- and time-dependent manner. Within the first 15-minutes, conduction velocity was unchanged in cell preparations treated with 10 μg/mL DEHP, but slowed by 14.6-18.3% in cells treated with higher concentrations (50 μg/mL DEHP: 0.215±0.06; 100 μg/mL DEHP: 0.225±0.04 mm/ms) relative to control cells (0.263±0.02 mm/ms; **Figure 8A,C**). With a longer 3 hour exposure, conduction velocity slowed by 12.9% in cells treated with 10 μg/mL DEHP (0.210±0.02 mm/ms) and 35.3-37% in cells treated with higher concentrations (50 DEHP: 0.156±0.05; 100 DEHP: 0.152±0.04 mm/ms) relative to control cells (0.241±0.02 mm/ms). Although conduction velocity measurements were comparable between 50 and 100 μg/mL DEHP, it became noticeably harder to pace hiPSC-CM at the higher DEHP concentration (**Figure 8B**). As such, the subsequent inhibitor studies were performed using only the intermediate dose of 50 μg/mL DEHP. Pretreatment with either atropine or doxycycline appeared to abrogate the DEHP-induced conduction velocity slowing, up to 1 hour of DEHP exposure (**Figure 8D,E**). However, with a longer 3-hour DEHP exposure time, hiPSC-CM conduction velocity slowed by 22.7% despite pretreatment with atropine (atropine: 0.229±0.02; 50 DEHP+atropine: 0.177±0.03 mm/ms) and 21.8% despite pretreatment with doxycycline (doxy: 0.243±0.02, 50 DEHP+doxy: 0.19±0.03 mm/ms).

**Figure 8.**
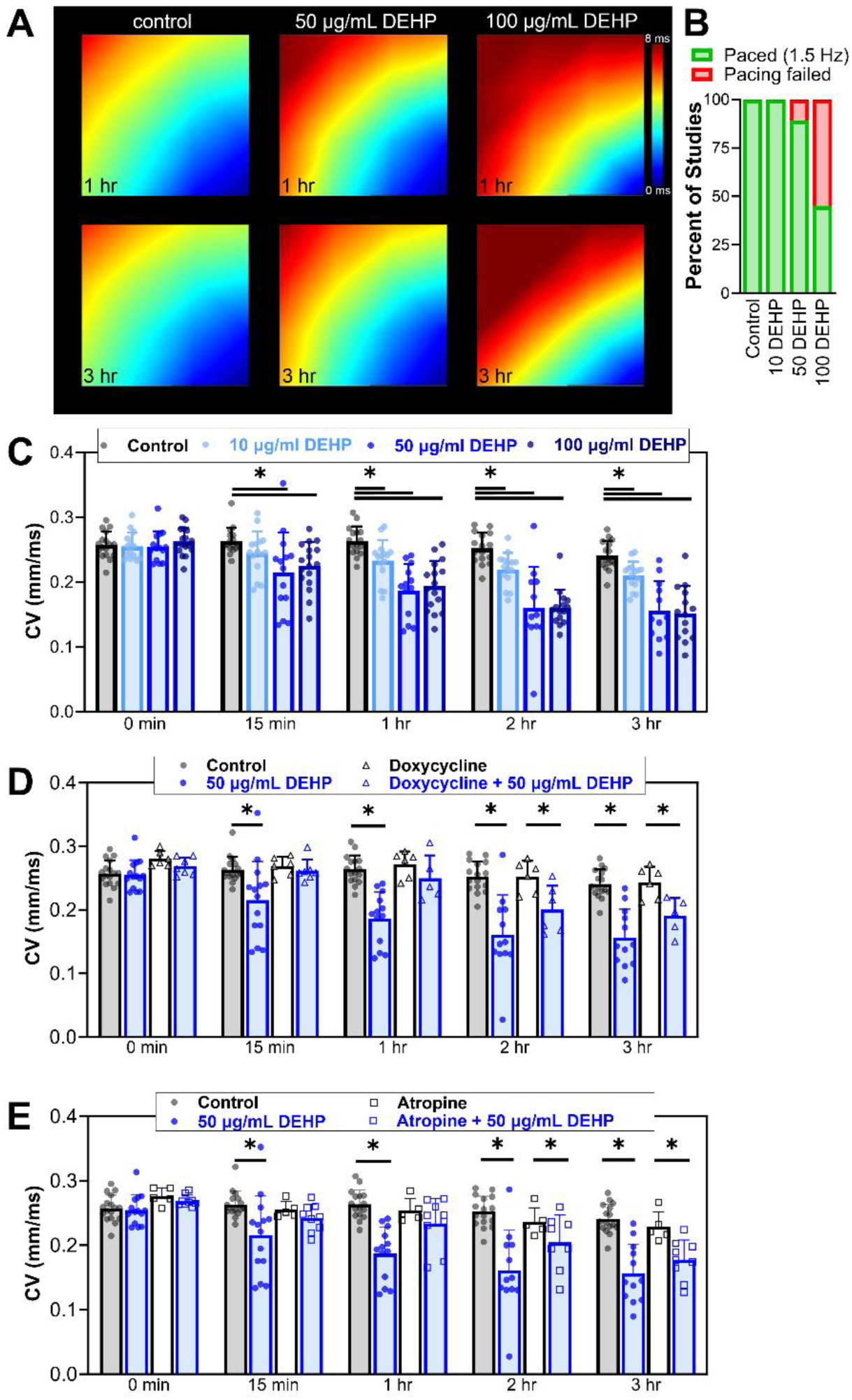
DEHP slows conduction velocity in hiPSC-CM monolayers. Conduction velocity was measured across hiPSC-CM monolayers plated atop microelectrode arrays, in response to external pacing (1.5 Hz). **A)** Activation maps show an electrical signal propagating from lower right corner (pacing electrode) across the hiPSC-CM monolayer after 1-3 hours treatment with control media or media supplemented with 50-100 μg/mL DEHP. **B)** hiPSC-CM treated with higher concentrations of DEHP were less likely to pace at the desired frequency and were omitted from analysis. **C)** Conduction velocity slowing in the presence of DEHP is time- and dose-dependent. **D,E)** After 15-60 minutes DEHP treatment, the effect on conduction velocity is reduced in the presence of either doxycycline (matrix metalloproteinase inhibitor) or atropine (muscarinic antagonist) - but DEHP-induced conduction velocity slowing persists after prolonged 2-3 hour treatment. *Comparisons were analyzed by two-way ANOVA with Holm-Sidak test for multiple comparisons. Significance between control and DEHP denoted by *p<0.05. n≥5 individual cell preparations per measurement*.

## DISCUSSION

Herein, we demonstrate that DEHP can exert cardiotoxic effects (slowed sinus rate, delayed sinus node recovery time, and prolonged AV conduction) in a dose- and time-dependent manner. Our findings build upon our previous work on MEHP(Jaimes, McCullough, Siegel, Swift, McInerney, *et al*., 2019), suggesting that both the parent chemical (DEHP) and its primary metabolite (MEHP) collectively perturb cardiac electrophysiology. The current study also builds upon the work of others, which report that low dose DEHP exposure (4 μg/ml) halts the spontaneous beating activity of chick cardiomyocytes within 30-minutes and elicits cell death within 24-hour exposure(Rubin and Jaeger, 1973). DEHP exposure has also been shown to precipitate bradycardia in the rat heart after 15-60 minutes(Aronson *et al*., 1978) – while *in vivo* MEHP exposure (primary metabolite of DEHP) results in cardiac arrest in a rodent model within minutes(Rock *et al*., 1987). In the current study, we also report that DEHP slows atrioventricular conduction, as measured by optical mapping, microelectrode array recordings, and electrocardiograms (PR interval) – wherein the later indicates AV conduction delay that progresses to heart block with longer exposure time (60-90 minutes). Previous work by Pfuderer et al. and Barry et al. suggested that phthalates exert cardiodepressive effects likely through interaction with cholinergic receptors, which can be rescued with atropine(Barry *et al*., 1990; Pfuderer and Francis, 1975). Although in our current study, pretreatment with 1 μM atropine did not prevent the effects of DEHP on either sinus rate or AV conduction.

Phthalate chemicals have also been shown to inhibit gap junction intercellular communication (GJIC) across multiple cell types(Gillum *et al*., 2009; Sobarzo *et al*., 2015, 2009; Cruciani *et al*., 1997; Isenberg *et al*., 2001), wherein the degree of inhibition is influenced by the length of the carbon chains(Čtveráčková *et al*., 2020). As an example, a comparative study conducted in liver cells reported that 80 μM DEHP reduced GJIC by ∼20% within 1 hour and ∼80% within 16 hours of exposure(Čtveráčková *et al*., 2020). Although a few reports have suggested that phthalate-disruption of intercellular coupling is species-dependent, at least in liver cells, wherein GJIC is impaired in mouse and rat hepatocytes, but not hamster, primate, or human hepatocytes(Isenberg *et al*., 2000; Kamendulis *et al*., 2002). Notably, in heart tissue, intercellular gap junction coupling is a key determinant of electrical conduction speed. In the current study, we report that 60-minute DEHP exposure slows epicardial conduction velocity by ∼15% in intact rat heart preparations. Further, we observed a similar effect in a human cardiomyocyte model (hiPSC-CM monolayer), wherein conduction velocity slowing was exaggerated with prolonged DEHP exposure (∼37% at 3 hours). To date, the mechanisms by which phthalate chemicals impair GJIC appear to be multifactorial. First, the lipophilic nature of DEHP (and other phthalates) facilitate its physical incorporation into cell membranes – as demonstrated by radiolabeling studies whereby DEHP localizes to the membrane fraction of RBCs(Rock *et al*., 1984). This physical disruption of cell membrane fluidity and organization has been confirmed by molecular dynamic simulation studies, which report that DEHP incorporates and aggregates in lipid bilayers(Bider *et al*., 2020). In this way, DEHP and other phthalates may act in a manner similar to long-chain alcohols or fatty acids to disrupt GJIC(Burt *et al*., 1991; Bastiaanse *et al*., 1993). Second, DEHP and other phthalates can disrupt the trafficking and/or stabilization of gap junction proteins at the cell membrane, by way of increased oxidative stress and matrix metalloproteinase activity(Yao *et al*., 2010; Sobarzo *et al*., 2015). Intercellular coupling is impaired under conditions of oxidative stress, due to the perturbation of forward trafficking and delivery of connexons to the cell membrane (Smyth *et al*., 2010). Intercellular coupling is also impaired with increased matrix metalloproteinase activity, as these enzymes can modulate electrical activity through the cleavage of connexin proteins(Lindsey *et al*., 2006; Wang *et al*., 2014). We found that pretreatment with an matrix metalloproteinase inhibitor (doxycycline) blocked the effects of DEHP on heart rate and sinus node recovery time in intact heart preparations, but AV conduction slowing persisted. Further, the use of an matrix metalloproteinase inhibitor reduced the effect of DEHP on conduction velocity in hiPSC-CM exposed for 15-60 minutes, but conduction velocity slowing persisted with longer exposure time (2-3 hours DEHP). Additional work is warranted to fully elucidate the mechanisms by which DEHP perturbs cardiac electrophysiology, and to clarify the degree to which its effects vary between different species and cell types.

DEHP has been widely used as a plasticizer additive in PVC-based medical products since the 1950s(Walter, 1984; Walter and MurphyY, 1952); nevertheless, several studies have documented DEHP leaching from plastic tubing circuits and intravenous or blood storage bags (D’alessandro *et al*., 2016; Rael *et al*., 2009; Loff *et al*., 2004, 2002, 2000; Karle *et al*., 1997; Inoue *et al*., 2005). As such, patients can incur phthalate exposure when undergoing multiple medical interventions that employ plastic products, including blood transfusions, cardiopulmonary bypass, and extracorporeal membrane oxygenation(Green *et al*., 2005; Weuve *et al*., 2006; Calafat *et al*., 2004; Mallow and Fox, 2014; Huygh *et al*., 2015; Kaestner *et al*., 2020; Eckert *et al*., 2020). Indeed, cardiac surgery patients often require blood transfusions due to priming of the cardiopulmonary bypass circuit, hemodilution and hemolysis during surgery, susceptibility to organ ischemia, blood loss, and low tolerance for anemia. Approximately 58-79% of all pediatric cardiac surgery patients receive at least one RBC transfusion(Mazine *et al*., 2015; Kato *et al*., 2020). Further, neonatal surgery patients (<5kg) receive blood products in massive quantity – equating to roughly 50% of the neonate’s total blood volume(Sturmer *et al*., 2018). These patients can be exposed to phthalate chemicals from both blood storage bags and DEHP-tubing used for cardiopulmonary bypass(Ramadan *et al*., 2020). Given the safety concerns associated with phthalate exposure, there is renewed interest in reducing the use of DEHP-containing plastics during blood donation, storage, and transfusion procedures(van der Meer *et al*., 2014; Prowse *et al*., 2014). Although, to date, the adoption of DEHP-free plastics has been limited due to availability, cost, and concern that alternative materials are inferior for blood storage applications(van der Meer *et al*., 2017; Razatos *et al*., 2022). Our study highlights the need for future work to investigate the impact of phthalate chemical exposure on vulnerable patient groups with heightened DEHP exposure. Notably, the negative effects of DEHP on cardiac electrophysiology occurred within 15-90 minutes, which is well within the timeframe of clinical exposure. Accordingly, our study is both clinically relevant and timely, as the results of this work may incentivize the development and adoption of DEHP-free materials for cardiac surgery, transfusion, and intensive care – or alternative mitigation strategies to reduce chemical exposure.

## LIMITATIONS

Since DEHP is a non-persistent chemical, the scope of our study was limited to acute DEHP exposure (15 min – 3 hours). An isolated, intact heart preparation was perfused with crystalloid buffer to optically measure cardiac action potentials and intracellular calcium transients – as blood perfusion interferes with fluorescent dye measurements. As such, the results of these *ex vivo* experiments may differ from the *in vivo* environment. Since *ex vivo* experiments were limited to a rodent model, differences with human physiology should be considered since prominent species-specific characteristics exist(Ripplinger *et al*., 2022). To address human applicability, we also employed an hiPSC-CM model for conduction velocity measurements; however, this simplified two-dimensional model lacks the complexity of the internal cardiac conduction system and prevents any electrical measurements between chambers of the heart (e.g., interatrial or atrioventricular conduction). hiPSC-CM are frequently used for preclinical studies of drug efficacy and safety, as these cells express the same ion channels that make up the human cardiac action potential. However, it is important to note that hiPSC-CM have an immature phenotype that more closely resembles a fetal-infant cardiomyocyte versus an adult cardiomyocyte(Lundy *et al*., 2013; Karbassi *et al*., 2020). Finally, the reported results are limited to the effect of DEHP, and therefore, do not take into account additional effects caused by its metabolites including MEHP.

## ACKNOWLEDGEMENTS

The authors acknowledge Blake Cooper for assistance with hiPSC-CM experiments, Anika Haski and Oluwatomisin Tiam Fotie for assistance with Langendorff-perfusion experiments, and Francesca Cendali and Daniel Stephenson PhD for assistance with phthalate chemical quantitation. We additionally acknowledge Dr. Meghan Delaney and Antoine Tavares Da Souza in the Children’s National Blood Bank for their assistance with procuring, storing, and collecting RBC units for this study.

## References

1. Aronson, C.E. et al. (1978) Effects of di-2-ethylhexyl phthalate on the isolated perfused rat heart. Toxicol Appl Pharmacol, 44, 155–169.

2. AuBuchon, J.P. et al. (1988) The effect of the plasticizer di-2-ethylhexyl phthalate on the survival of stored RBCs. Blood, 71, 448–52.

3. Bagel-Boithias, S. et al. (2005) Leaching of diethylhexyl phthalate from multilayer tubing into etoposide infusion solutions. Am J Health Syst Pharm, 62, 182–188.

4. Barry, Y.A. et al. (1990) Atropine inhibition of the cardiodepressive effect of mono(2-ethylhexyl)phthalate on human myocardium. Toxicol Appl Pharmacol, 106, 48–52.

5. Bastiaanse, E.M.L. et al. (1993) Heptanol-induced decrease in cardiac gap junctional conductance is mediated by a decrease in the fluidity of membranous cholesterol-rich domains. J Membr Biol, 136, 135–145.

6. Bider, R.-C. et al. (2020) Stabilization of Lipid Membranes through Partitioning of the Blood Bag Plasticizer Di-2-ethylhexyl phthalate (DEHP).

7. Braun, J.M. et al. (2013) Phthalate exposure and children’s health Curr Opin Pediatr, Department of Epidemiology.

8. Burt, J.M. et al. (1991) Uncoupling of cardiac cells by fatty acids: structure-activity relationships. 10.1152/ajpcell.1991.260.3.C439, 260.

9. Calafat, A.M. et al. (2004) Exposure to di-(2-ethylhexyl) phthalate among premature neonates in a neonatal intensive care unit. Pediatrics, 113, e429–34.

10. Cooper, B.L. et al. (2021) KairoSight: Open-Source Software for the Analysis of Cardiac Optical Data Collected From Multiple Species. Front Physiol, 0, 1820.

11. Cruciani, V. et al. (1997) Effects of Peroxisome Proliferators and 12-O-tetradecanoyl phorbol-13-acetate on Intercellular Communication and connexin43 in Two Hamster Fibroblast Systems. Int J Cancer, 73.

12. Čtveráčková, L. et al. (2020) Structure-Dependent Effects of Phthalates on Intercellular and Intracellular Communication in Liver Oval Cells. Int J Mol Sci, 21, 1–21.

13. D’alessandro, A. et al. (2016) Rapid detection of DEHP in packed red blood cells stored under European and US standard conditions. Blood Transfus, 14, 140–4.

14. DiGangi, J. (1999) Phthalates in PVC medical products from 12 countries Greenpeace.

15. Eckert, E. et al. (2020) Plasticizer exposure of infants during cardiac surgery. Toxicol Lett, 330, 7–13.

16. Fedorov, V. V. et al. (2007) Application of blebbistatin as an excitation-contraction uncoupler for electrophysiologic study of rat and rabbit hearts. Heart rhythm : the official journal of the Heart Rhythm Society, 4, 619–626.

17. Gillum, N. et al. (2009) Clinically relevant concentrations of di (2-ethylhexyl) phthalate (DEHP) uncouple cardiac syncytium. Toxicol Appl Pharmacol, 236, 25–38.

18. Glynn, S.A. et al. (2016) The red blood cell storage lesion: the end of the beginning. Transfusion (Paris), 56, 1462–1468.

19. Green, R. et al. (2005) Use of di(2-ethylhexyl) phthalate-containing medical products and urinary levels of mono(2-ethylhexyl) phthalate in neonatal intensive care unit infants. Environ Health Perspect, 113, 1222–1225.

20. Haq, K.T. et al. (2023) KairoSight-3.0 : A Validated Optical Mapping Software to Characterize Cardiac Electrophysiology, Excitation-Contraction Coupling, and Alternans. bioRxiv.

21. Hoover, D.B. and Neely, D.A. (1997) Differentiation of Muscarinic Receptors Mediating Negative Chronotropic and Vasoconstrictor Responses to Acetylcholine in Isolated Rat Hearts. Journal of Pharmacology and Experimental Therapeutics, 282.

22. Horowitz, B. et al. (1985) Stabilization of red blood cells by the plasticizer, diethylhexylphthalate. Vox Sang, 48, 150–5.

23. Huygh, J. et al. (2015) Considerable exposure to the endocrine disrupting chemicals phthalates and bisphenol-A in intensive care unit (ICU) patients. Environ Int, 81, 64–72.

24. Inoue, K. et al. (2005) Evaluation and analysis of exposure levels of di(2-ethylhexyl) phthalate from blood bags. Clin Chim Acta, 358, 159–166.

25. Isenberg, J.S. et al. (2000) Effects of Di-2-Ethylhexyl Phthalate (DEHP) on Gap-Junctional Intercellular Communication (GJIC), DNA Synthesis, and Peroxisomal Beta Oxidation (PBOX) in Rat, Mouse, and Hamster Liver. Toxicological Sciences, 56, 73–85.

26. Isenberg, J.S. et al. (2001) Reversibility and persistence of di-2-ethylhexyl phthalate (DEHP)- and phenobarbital-induced hepatocellular changes in rodents. Toxicol Sci, 64, 192–199.

27. Jaeger, R.J. and Rubin, R.J. (1973) Extraction, localization, and metabolism of di-2-ethylhexyl phthalate from PVC plastic medical devices. Environ Health Perspect, 3, 95–102.

28. Jaimes, R. et al. (2016) A technical review of optical mapping of intracellular calcium within myocardial tissue. American Journal of Physiology-Heart and Circulatory Physiology, 310, H1388–401.

29. Jaimes, R., McCullough, D., Siegel, B., Swift, L., Hiebert, J., et al. (2019) Lights, camera, path splitter: a new approach for truly simultaneous dual optical mapping of the heart with a single camera. BMC Biomed Eng, 1, 25.

30. Jaimes, R., McCullough, D., Siegel, B., Swift, L., McInerney, D., et al. (2019) Plasticizer Interaction with the Heart: Chemicals Used in Plastic Medical Devices Can Interfere with Cardiac Electrophysiology. Circ Arrhythm Electrophysiol, 12.

31. Kaestner, F. et al. (2020) Exposure of patients to di(2-ethylhexy)phthalate (DEHP) and its metabolite MEHP during extracorporeal membrane oxygenation (ECMO) therapy. 15, e0224931.

32. Kamendulis, L.M. et al. (2002) Comparative effects of phthalate monoesters on gap junctional intercellular communication and peroxisome proliferation in rodent and primate hepatocytes. J Toxicol Environ Health A, 65, 569–588.

33. Karbassi, E. et al. (2020) Cardiomyocyte maturation: advances in knowledge and implications for regenerative medicine. Nat Rev Cardiol.

34. Karle, V.A. et al. (1997) Extracorporeal membrane oxygenation exposes infants to the plasticizer, di(2-ethylhexyl)phthalate. Crit Care Med, 25, 696–703.

35. Kato, H. et al. (2020) Are Blood Products Routinely Required in Pediatric Heart Surgery? Pediatr Cardiol, 41, 932–938.

36. Labow, R.S. et al. (1987) The effect of the plasticizer di(2-ethylhexyl)phthalate on red cell deformability. Blood, 70, 319–23.

37. Lindsey, M.L. et al. (2006) Matrix metalloproteinase-7 affects connexin-43 levels, electrical conduction, and survival after myocardial infarction. Circulation, 113, 2919–2928.

38. Loff, S. et al. (2004) Extraction of di-ethylhexyl-phthalate from perfusion lines of various material, length and brand by lipid emulsions. J Pediatr Gastroenterol Nutr, 39, 341–345.

39. Loff, S. et al. (2002) Kinetics of diethylhexyl-phthalate extraction From polyvinylchloride-infusion lines. JPEN J Parenter Enteral Nutr, 26, 305–309.

40. Loff, S. et al. (2000) Polyvinylchloride infusion lines expose infants to large amounts of toxic plasticizers. J Pediatr Surg, 35, 1775–1781.

41. Di Lorenzo, M. et al. (2020) Intrauterine exposure to diethylhexyl phthalate disrupts gap junctions in the fetal rat testis. Curr Res Toxicol, 1, 5–11.

42. Lundy, S.D. et al. (2013) Structural and functional maturation of cardiomyocytes derived from human pluripotent stem cells. Stem Cells Dev, 22, 1991–2002.

43. Malarvannan, G. et al. (2019) Phthalate and alternative plasticizers in indwelling medical devices in pediatric intensive care units. J Hazard Mater, 363, 64–72.

44. Mallow, E.B. and Fox, M.A. (2014) Phthalates and critically ill neonates: device-related exposures and non-endocrine toxic risks. Journal of Perinatology, 34, 892–897.

45. Mazine, A. et al. (2015) Blood Transfusions After Pediatric Cardiac Operations: A North American Multicenter Prospective Study. Ann Thorac Surg, 100, 671–677.

46. van der Meer, P.F. et al. (2017) Alternatives in blood operations when choosing non-DEHP bags. Vox Sang, 112, 183–184.

47. van der Meer, P.F. et al. (2014) Should DEHP be eliminated in blood bags? Vox Sang, 106, 176–195.

48. Nemkov, T. et al. (2019) High-throughput metabolomics: Isocratic and gradient mass spectrometry-based methods. In, Methods in Molecular Biology. Humana Press Inc., pp. 13–26.

49. Pfuderer, P. and Francis, A.A. (1975) Phthalate esters: Heartrate depressors in the goldfish. Bull Environ Contam Toxicol, 13, 275–279.

50. Posnack, N.G. et al. (2015) Exposure to phthalates affects calcium handling and intercellular connectivity of human stem cell-derived cardiomyocytes. PLoS One, 10, e0121927–e0121927.

51. Posnack, N.G. et al. (2011) Gene expression profiling of DEHP-treated cardiomyocytes reveals potential causes of phthalate arrhythmogenicity. Toxicology, 279, 54.

52. Posnack, N.G. (2021) Plastics and cardiovascular disease. Nat Rev Cardiol, 18, 69–70.

53. Posnack, N.G. (2014) The Adverse Cardiac Effects of Di(2-ethylhexyl)phthalate and Bisphenol A. Cardiovasc Toxicol, 14, 339–357.

54. Prowse, C. V. et al. (2014) Commercially available blood storage containers. Vox Sang, 106, 1–13.

55. Rael, L.T. et al. (2009) Phthalate esters used as plasticizers in packed red blood cell storage bags may lead to progressive toxin exposure and the release of pro-inflammatory cytokines. Oxid Med Cell Longev, 2, 166–71.

56. Ramadan, M. et al. (2020) Bisphenols and phthalates: Plastic chemical exposures can contribute to adverse cardiovascular health outcomes. Birth Defects Res.

57. Razatos, A. et al. (2022) Survey of blood centre readiness regarding removal of DEHP from blood bag sets: The BEST Collaborative Study. Vox Sang, 117, 796–802.

58. Ripplinger, C.M. et al. (2022) Guidelines for assessment of cardiac electrophysiology and arrhythmias in small animals. Am J Physiol Heart Circ Physiol, 323, H1137–H1166.

59. Rock, G. et al. (1986) Distribution of di(2-ethylhexyl) phthalate and products in blood and blood components. Environ Health Perspect, 65, 309.

60. Rock, G. et al. (1987) Hypotension and cardiac arrest in rats after infusion of mono(2-ethylhexyl) phthalate (MEHP), a contaminant of stored blood. N Engl J Med, 316, 1218–9.

61. Rock, G. et al. (1984) Incorporation of plasticizer into red cells during storage. Transfusion (Paris), 24, 493–498.

62. Rose, R.J. et al. (2012) The effect of temperature on di(2-ethylhexyl) phthalate leaching from PVC infusion sets exposed to lipid emulsions. Anaesthesia, 67, 514–520.

63. Rubin, R.J. and Jaeger, R.J. (1973) Some pharmacologic and toxicologic effects of di-2-ethylhexyl phthalate (DEHP) and other plasticizers. Environ Health Perspect, 3, 53–59.

64. Scannevin, R.H. et al. (2017) Discovery of a highly selective chemical inhibitor of matrix metalloproteinase-9 (MMP-9) that allosterically inhibits zymogen activation. Journal of Biological Chemistry, 292, 17963–17974.

65. Shang, J. et al. (2019) Recovery From a Myocardial Infarction Is Impaired in Male C57bl/6 N Mice Acutely Exposed to the Bisphenols and Phthalates That Escape From Medical Devices Used in Cardiac Surgery. Toxicol Sci, 168, 78–94.

66. Shelby, M.D. (2006) NTP-CERHR monograph on the potential human reproductive and developmental effects of di (2-ethylhexyl) phthalate (DEHP). NTP CERHR MON, v, vii–7, II-iii-xiii passim.

67. Sjoberg, P.O. et al. (1985) Exposure of newborn infants to plasticizers. Plasma levels of di-(2-ethylhexyl) phthalate and mono-(2-ethylhexyl) phthalate during exchange transfusion. Transfusion (Paris), 25, 424–428.

68. Smyth, J.W. et al. (2010) Limited forward trafficking of connexin 43 reduces cell-cell coupling in stressed human and mouse myocardium. 120, 266–279.

69. Sobarzo, C.M. et al. (2009) Effects of di(2-ethylhexyl) phthalate on gap and tight junction protein expression in the testis of prepubertal rats. Microsc Res Tech, 72, 868–877.

70. Sobarzo, C.M. et al. (2015) Mono-(2-ethylhexyl) phthalate (MEHP) affects intercellular junctions of Sertoli cell: A potential role of oxidative stress. Reprod Toxicol, 58, 203–212.

71. Sturmer, D. et al. (2018) Recent innovations in perfusion and cardiopulmonary bypass for neonatal and infant cardiac surgery. Transl Pediatr, 7, 139–150.

72. Swift, L.M. et al. (2019) Optocardiography and Electrophysiology Studies of Ex Vivo Langendorff-perfused Hearts.

73. Swift, L.M. et al. (2012) Properties of blebbistatin for cardiac optical mapping and other imaging applications. Pflugers Arch Eur J Physiol, 464, 503–512.

74. Swift, L.M. et al. (2021) Stop the beat to see the rhythm: excitation-contraction uncoupling in cardiac research. Am J Physiol Heart Circ Physiol, 321, H1005–H1013.

75. Takahashi, Y. et al. (2008) Di(2-ethylhexyl) phthalate exposure during cardiopulmonary bypass. Asian Cardiovasc Thorac Ann, 16, 4–6.

76. Tereshchenko, L.G. and Posnack, N.G. (2019) Does plastic chemical exposure contribute to sudden death of patients on dialysis? Heart Rhythm, 16, 312–317.

77. Trasande, L. et al. (2022) Phthalates and attributable mortality: A population-based longitudinal cohort study and cost analysis. Environ Pollut, 292, 118021.

78. Walter, C.W. (1984) Invention and development of the blood bag. Vox Sang, 47, 318–324.

79. Walter, C.W. and MurphyY, W.P. (1952) A closed gravity technique for the preservation of whole blood in ACD solution utilizing plastic equipment. Surg Gynecol Obstet, 94, 687–692.

80. Wang, J. et al. (2014) Dynamic alterations of connexin43, matrix metalloproteinase-2 and tissue inhibitor of matrix metalloproteinase-2 during ventricular fibrillation in canine. Mol Cell Biochem, 391, 259–266.

81. Weuve, J. et al. (2006) Exposure to phthalates in neonatal intensive care unit infants: urinary concentrations of monoesters and oxidative metabolites. Environ Health Perspect, 114, 1424–1431.

82. Yao, P.-L. et al. (2010) Mono-(2-Ethylhexyl) Phthalate-Induced Disruption of Junctional Complexes in the Seminiferous Epithelium of the Rodent Testis Is Mediated by MMP21. Biol Reprod, 82, 516–527.

83. Zeng, G. et al. (2022) Low-level plasticizer exposure and all-cause and cardiovascular disease mortality in the general population. Environ Health, 21, 1–11.

84. Zhao, Y.X. et al. (2022) Gap Junction Protein Connexin 43 as a Target Is Internalized in Astrocyte Neurotoxicity Caused by Di-(2-ethylhexyl) Phthalate. J Agric Food Chem, 70, 5921–5931.

